# Reproductive outcomes in sport horse commercial French breeding farms: clinical relevance of mares’ age, parity and lactating status

**DOI:** 10.1101/2022.01.21.477204

**Authors:** Emilie Derisoud, Bénédicte Grimard, Clothilde Gourtay, Juliette Auclair-Ronzaud, Pascale Chavatte-Palmer

**Affiliations:** Department of Physiology and Pharmacology, Karolinska Institutet, Solna, Stockholm, Sweden; Université Paris-Saclay, UVSQ, INRAE, BREED, 78350, Jouy-en-Josas, France; Ecole Nationale Vétérinaire d’Alfort, BREED, 94700, Maisons-Alfort, France; IFCE, Plateau technique du Pin, 61310, Exmes, France; IFCE, Plateau technique de Chamberet, 19370, Chamberet, France

**Author notes:** authors contributed equally to this work (Emilie Derisoud, Bénédicte Grimard). Mail address: Pascale Chavatte-Palmer, BREED (UMR1198) – Bâtiment 230, Biology de la reproduction, Environnement, Epigénétique et Développement Domaine de Vilvert, 78352 Jouy-en-Josas, FRANCE.

**Keywords:** Horse, equine, reproduction, insemination, lactation, fertility

## Abstract

**Background:** For a long time, important and progressive fertility decline is observed with mare age. Parity and yearly breeding, however, are controversial factors impacting fertility. Most of these studies were performed on race horses and before 2010, on data accumulated before. The cumulative effects of age and nulliparity/nursing with improved reproductive technologies and practices, often observed in sport horses, has not been investigated yet.

**Objectives:** To investigate the effect of age, parity and nursing, as well as reproductive management, on post-insemination inflammation and fertility in sports mares.

**Study design:** Age, parity, nursing status, follicular size before ovulation, estrus and/or ovulation induction, artificial insemination (AI) protocol, post-breeding inflammation and treatment, Day 14 pregnancies rates (PR) and number of embryos, as well as subsequent foaling the next year were recorded for 277 mares (506 cycles) over one breeding season in two French commercial studs.

**Methods:** Multivariate logistic regression models were used.

**Results:** PR was 41.9% per cycle and 76.5% at the end of the season. Post-insemination fluid accumulation risk was increased in >10-year-old, barren or July or August inseminated mares (p<0.0001, OR=3.29, 5.389 and 3.329, respectively). PR were reduced in mares >10-year-old vs younger or inseminated with frozen vs fresh or refrigerated semen (p<0.05, OR=0.622 and 0.582, respectively). More pregnancy on Day 14 were observed in mares with multiple ovulations compared to mono- ovulation (p<0.05, OR=1.791). Regardless the age, PR only tended to be improved in multiparous (p=0.07, OR=1.434 in parous vs nulliparous) but in >10-year-old mares, multiparity greatly increased PR (44.09% in parous vs 30.89% in nulliparous, p<0.05).

**Main limitations:** Limited number of mares.

**Conclusion:** In sport horse, maternal age was more important than parity and lactating status for reproductive success but cumulative effects of nulliparity and aging was deleterious on PR, demonstrating their importance in the management of sport mares.

## 1. Introduction

Effectiveness of mare reproduction is an important economical factor in the horse industry. Once mares begin their reproductive career, breeders generally aim to produce one foal per year per mare to remain profitable. This goal can only be achieved if mares remain fertile throughout their life span.

The decline in mare fertility associated with age is well described [1–18] but mainly on Thoroughbreds where the career is short and mares are bred from and culled at younger age compared to sport horses. It is common in sport horses to breed older mares: in Canada, 37% of the standardbred mares [19] and in Finland, 53% of Standardbred and 63% of Finnhorse mares exceed 11 years old at insemination [3]. Furthermore, mares can be bred shortly after foaling and are often nursing at the time of breeding. Lactation affects mares’ metabolism [20] but there is no consensus on whether it may induce effects on fertility and embryo/fetal loss [21–24]. As for lactation, the putative effects of mare’s parity on fertility and embryo loss are controversial [2–5,9,10,12–16,18]. Plus, in sport horses, mares are often both nulliparous and old at the same time and in Finland, 20.5% of Finnhorse and 15.5% of Standardbred broodmares are both older than 10 and nulliparous [3].

Transient uterine inflammation is a normal physiological reaction in mares after breeding. For some mares, however, a persistent infection can develop and interfere with fertility outcomes (for review [25]). Indeed, 45% (N=22) of the 8-16 years old mares and 88% of the older mares (≥17 years old, N=26) had uterine fluid retention 48h after insemination with frozen semen, whereas none of the 9 younger mares was affected [26]. Parity and/or lactation on the post-breeding inflammation have not been explored yet.

The Thoroughbred studbook only allows hand breeding whereas the French Trotter studbook allows only insemination with fresh semen. Such limitations do not exist in sport horse and no recent study considered at the same time the method of reproduction and the effects of age, parity and lactation on mare fertility.

This study aimed to identify effects of mare age, parity and nursing on post-breeding fluid accumulation, fertility and embryonic/fetal death in commercial stud conditions for sport mares.

## 2. Material and methods

### 2.1. Animals

Data were recorded in 2 French commercial stud farms, one located about 60km south west of Paris, the other 150 km south of Paris, both managed by one artificial insemination cooperative during one breeding season (2019).

From beginning of February to late August 2019, the study enrolled 277 sport mares over 506 estrus periods. For each mare, date of birth, breed, number of previous breedings and foalings were extracted from the national database (Infochevaux: https://infochevaux.ifce.fr/fr/info-chevaux). Mares were clustered according to age in two (≤ 10 years old *vs* > 10 years old) or four classes (≤ 5 years old, ]5, 10], ]10, 15], > to 15 years old). They were also clustered in 2 classes according to previous fertility (number of previous foaling/number of previous breedings <0.7 or >0.7) and in 2 classes according to parity: nulliparous (never foaled before) vs parous (foaled at least once before) at the beginning of the breeding season.

### 2.2. Breeding management

#### 2.2.1. Estrus monitoring

First, luteolysis was induced in mares that had a persistent corpus luteum or that were sent for breeding late in the season, using prostaglandin F2α analogs: Alfaprostol (Alfabedyl©) or Luprostol (Prosolvin®, 1 to 1.5 mL IM according to the mare size).

Estrus detection was performed using uterine horn firmness estimated by rectal palpation together with ultrasound examination where the uterine oedema score (0-5) [27] and the diameter of the largest follicle were determined. Mares were monitored once or twice daily (according to breeding method, see below) from the time when the diameter of the largest follicle reached 35 mm and until ovulation was detected.

As ovulation approached (follicle diameter ≥35 mm and reduction of the uterine oedema score), ovulation was induced with hCG (Chorulon®, 1500 to 5000 IU IM related to the mare weight) or with a GnRH analog (Decapeptyl®, 0.1 mg triptorelin in 1 mL SC). The GnRH analog option was chosen when mares had a known resistance to hCG or when hCG had already been used before during the breeding season and was not effective to induce ovulation.

#### 2.2.2. Insemination management

All mares were bred using artificial insemination (AI). Mares were inseminated with fresh (FAI), cooled transported (CTAI) or frozen semen (FZAI). As this study has been performed in a commercial breeding farm, insemination dose was dependent on the number of straws bought by the broodmare’s owner and the country where straws have been produced. Mare management differed according to the type of semen used.

Mares entitled to be inseminated with fresh semen were examined once daily during estrus. A first insemination was performed maximum 24h before ovulation (follicle diameter ≥ 35 mm and decrease in the uterine oedema score) and mares that had not ovulated 48h later were inseminated a second time. This procedure was repeated until ovulation was observed.

Mares entitled to be inseminated with cooled transported semen were also examined once daily and inseminated before ovulation, but subsequent inseminations were performed every 24h until ovulation.

Mares bred with frozen semen were examined every 12h if the number of available straws was >4 while examinations were performed every 6 h when the number of available straws was ≤4. In this latter case, deep horn insemination was performed. The aim was to inseminate as close as possible of ovulation (*i.e.* maximum 6 hours before or after ovulation), as determined by ultrasound observation of preovulatory follicular wall thickening and follicle deformation.

#### 2.2.3. Post-breeding fluid accumulation and pregnancy diagnosis

All mares were examined by ultrasound the day after insemination. If uterine oedema and/or fluid accumulation were observed, mares were treated either with a single dose of oxytocin (Ocytocine S©, 10-20 UI in 1 to 1.5 mL IM or IV), or oxytocin in association with uterine lavage (one or two litters of warm, sterile saline solution), or in association with antibiotics (in accordance with the recommendation of the use of antibiotics in veterinary medicine) or by uterine lavage alone. Treatment was decided by the veterinary surgeon and depended on the volume of fluid accumulated.

Pregnancy was assessed 14 days (D14) after AI by ultrasonography. In the case of twin pregnancy, squeezing was recommended to the breeder but not always performed. Pregnancy was confirmed on Day 30 if the mare was brought back to the stud. In addition, some mares came back again in autumn for late pregnancy diagnosis.

#### 2.2.4. Data recording

All data were recorded in the same dedicated software (Gynebase©, Equidéclic, France) by the veterinarians and technicians of the 2 studs.

Age, parity, lactational status at breeding (nursing *vs* non nursing), number of monitored cycles for each mare, estrus induction (yes/no), heat duration (number of days of observation by ultrasonography), size of the preovulatory follicle, induction of ovulation (yes/no), hormone used to induce ovulation (hCG or GnRH analog), ovulation observed (yes/no), number of follicles ovulated, AI mode (fresh semen, cooled transported or frozen semen), number of AI during estrus, date of insemination (last AI if more than one AI were performed during estrus), stallion identity, number of straws used (for frozen semen), uterine fluid accumulation after AI (yes/no), treatment used (oxytocin alone vs other), presence of single or twin embryos, squeezing when twins were diagnosed (yes/no), pregnancy diagnosis on D14 as well as on D30 or later, when possible, were recorded. The number of the monitored cycles needed to obtain a gestation was classified as 1 *vs* >1 when pregnancy was achieved after more than one cycle. The number of straws used for frozen insemination was classified as either <8 *vs* ≥ 8 or ≤ 4 vs > 4. Cutoff values of 4 and 8 were chosen according to previous (8 straws for 400 million mobile spermatozoa) and the current (4 straws of 50 x 10^6^ mobile spermatozoa, 4 mL each) recommendation of the French Institute for Horses and Horse-riding (IFCE) for frozen semen.

Pregnancy rate was calculated as the ratio between the number of pregnant mares at the end of the breeding season and total number of mares bred (overall pregnancy rate at the end of the season) and as the ratio of number of positive diagnoses on D14 / total number of used cycles (pregnancy rate per cycle). Embryonic loss was estimated by the number of mares not pregnant on D30 after a positive pregnancy diagnosis on D14. Late embryonic loss was calculated by the number of mares not pregnant during the autumn after a positive pregnancy diagnosis on D14. Foal birth was checked the next year using the national database. Total embryonic/fetal loss was defined as the number of mares that did not foal after a positive pregnancy diagnosis on D14. Mares were only considered as non-foaling when abortion/stillbirth or new breeding without foaling were recorded in the national database in 2020.

### 2.3. Statistical analysis

Data were analyzed using SAS® Studio 3.8 (SAS® University Edition).

#### 2.3.1. Univariate & multivariate analysis

Univariate analysis was used to evaluate the effects of qualitative variables on (1) the incidence of post-breeding fluid accumulation, on (2) pregnancy rate and (3) embryonic/fetal loss. For (1) and (2), all recorded cycles were used while for (3), only pregnant mares on D14 were considered. The statistical unit was, therefore, the cycle for (1) and (2) while it was the mare for (3).

For each criterion analyzed, results were compared between the different classes using a Chi-Squared test. Variables associated at p <0.20 were included in the second step of the analysis.

As a second step, multivariate analysis was conducted using logistic regression (GLIMMIX procedure of SAS® Studio). Individual effects were considered by including mare identity in each model as a random effect. A backward stepwise elimination of non-associated (p>0.10) variables was performed to develop multivariate models. Models presenting the lowest Akaike’s Information Criterion were retained.

For these analyses, breeds and stallions could not be used as their numbers were too high without enough repeats to be considered in the statistical analysis.

For quantitative data, results are presented as mean ± standard error.

#### 2.3.2. Analysis of age, parity and nursing status

The analyses were performed using the estrous cycle as the reference unit. Young (≤ 10 years) *vs* old (>10 years), nulliparous *vs* parous, nursing *vs* non nursing mares were compared using T test for quantitative variables and Xi Square for qualitative ones. For further characterization of the suckling status and mares’ parity, these factors were analyzed using Xi Square analysis in each population of young or old mares.

## 3. Results

### 3.1. Descriptive analysis

#### 3.1.1. Population of study

The 277 mares belonged to 32 breeds approved by the French Institute of the Horse (IFCE), with French saddlebred (Selle français) being the most represented breed (56% of mares included in this study). No other breed reached more than 15 individuals. Eight mares did not belong to any French approved breed. In addition, 13 mares belonged to racehorse breeds (3 English Thoroughbreds, 10 French Trotters). There were also 22 ponies and one draft horse. All mares were used for sport and/or leisure.

Characteristics of the 277 mares are detailed in Table 1. The average age was 12.7 years old (min 2, max 23, σ = 4.8). Altogether, 196 mares had been bred at least in one previous breeding season and the overall average of previous breeding seasons was 2.5 (min 0, max 17, σ = 3.0). For previously bred mares, mean fertility (number of foalings/number of breeding) was 0.71 (min 0, max 1, σ = 0.31). The interval between previous breeding and the 2019 reproductive season ranged from 1 to 13 years. Altogether, 66 (21.9%) mares were nulliparous and aged more than 10 years and 44 (15.8%) were suckling and older than 10 years.

**Table 1:** Characteristics of the 277 mares bred in 2019.

| Variable | n | Percent |
| --- | --- | --- |
| Age in 2 classes (Years) |  |  |
| $\leq 10$ | 96 | 34.7 |
| $> 10$ | 181 | 65.3 |
| Age in 4 classes (Years) |  |  |
| ≤5 | 24 | 8.7 |
| 5-9 | 72 | 26.0 |
| 10-14 | 94 | 33.9 |
| > 15 | 87 | 31.4 |
| Parity |  |  |
| Nulliparous | 98 | 35.4 |
| Parous | 179 | 64.6 |
| Lactation status in 2019 |  |  |
| Suckling | 76 | 27.4 |
| Non suckling | 201 | 72.6 |
| Fertility at previous breeding (n=196) |  |  |
| < 0.7 | 82 | 41.8 |
| ≥ 0.7 | 114 | 58.2 |
| Month of first AI |  |  |
| March | 36 | 13.0 |
| April | 70 | 25.3 |
| May | 86 | 31.0 |
| June | 67 | 24.2 |
| July - August | 18 | 6.5 |

#### 3.1.2. Pregnancy rate per mare

Mares were bred with 154 different stallions. Altogether, 212 mares were diagnosed pregnant on D14 at the end of the season (76.5% of D14 pregnancy). These performances were obtained within an average of 1.8 cycle per mare (min 1, max 6). From the 154 mares that came back for pregnancy confirmation, 130 were still pregnant at Day 30, which represents 9.1% of pregnancy loss. Only forty-six mares came back later in the autumn, of which 44 mares were confirmed still pregnant (4.3% of pregnancy loss during this period).

Foaling information in 2020 were obtained for 206 pregnant mares on Day 14. Among them, 168 foalings were recorded indicating a total embryonic/fetal loss of 18.4%.

#### 3.1.3. Breeding management

Data recorded per estrus period are presented on Table 2.

**Table 2:**
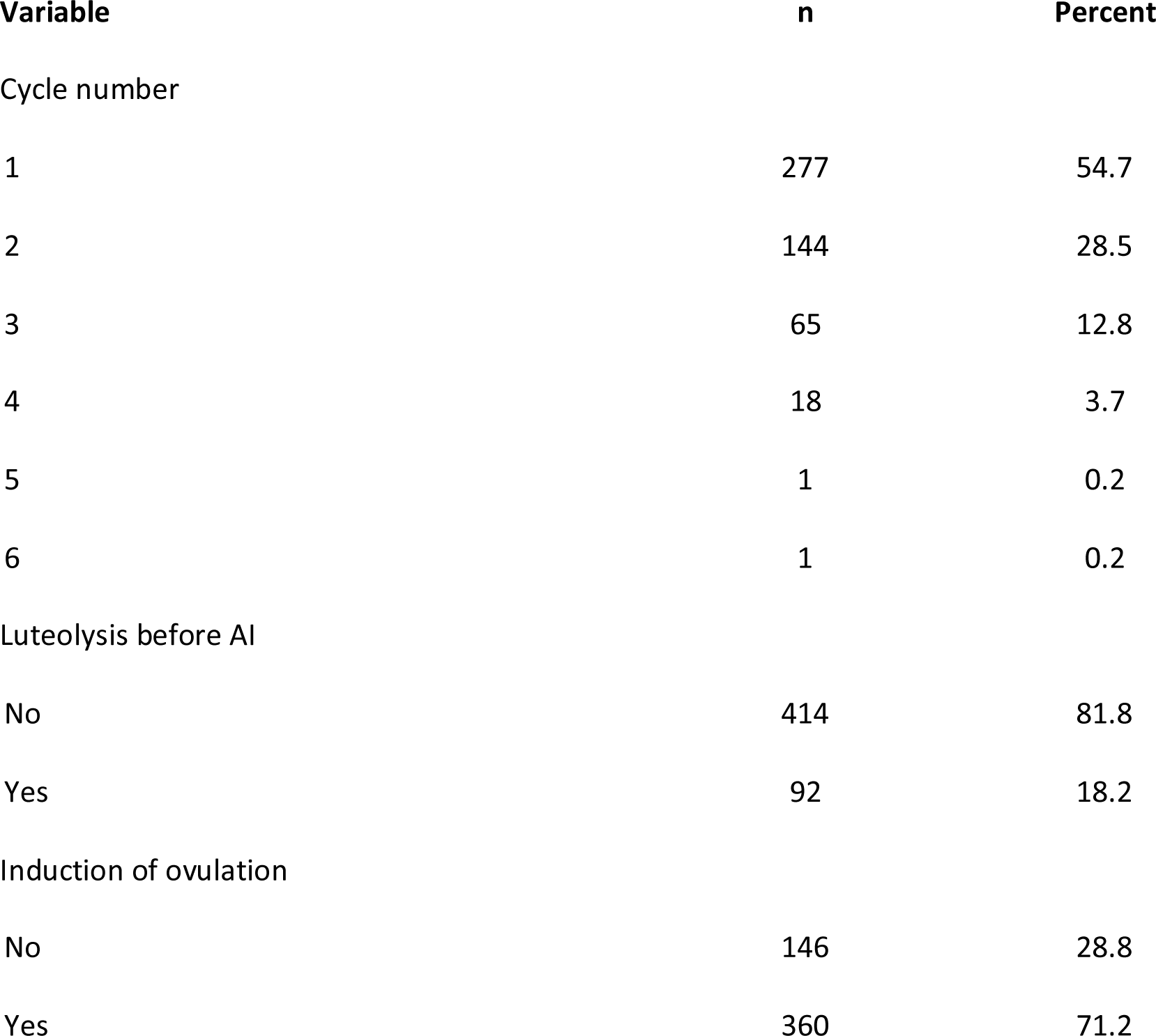
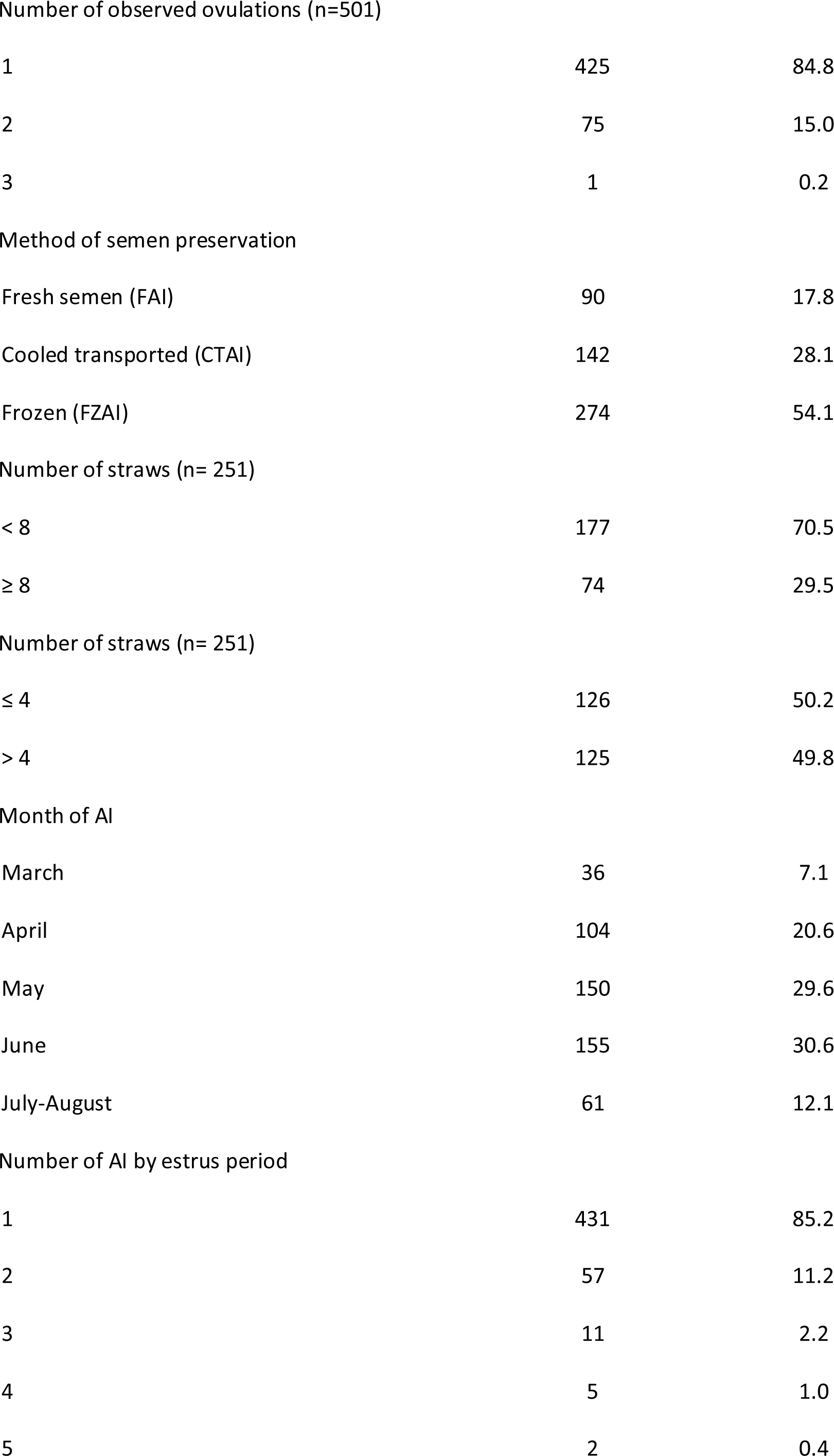
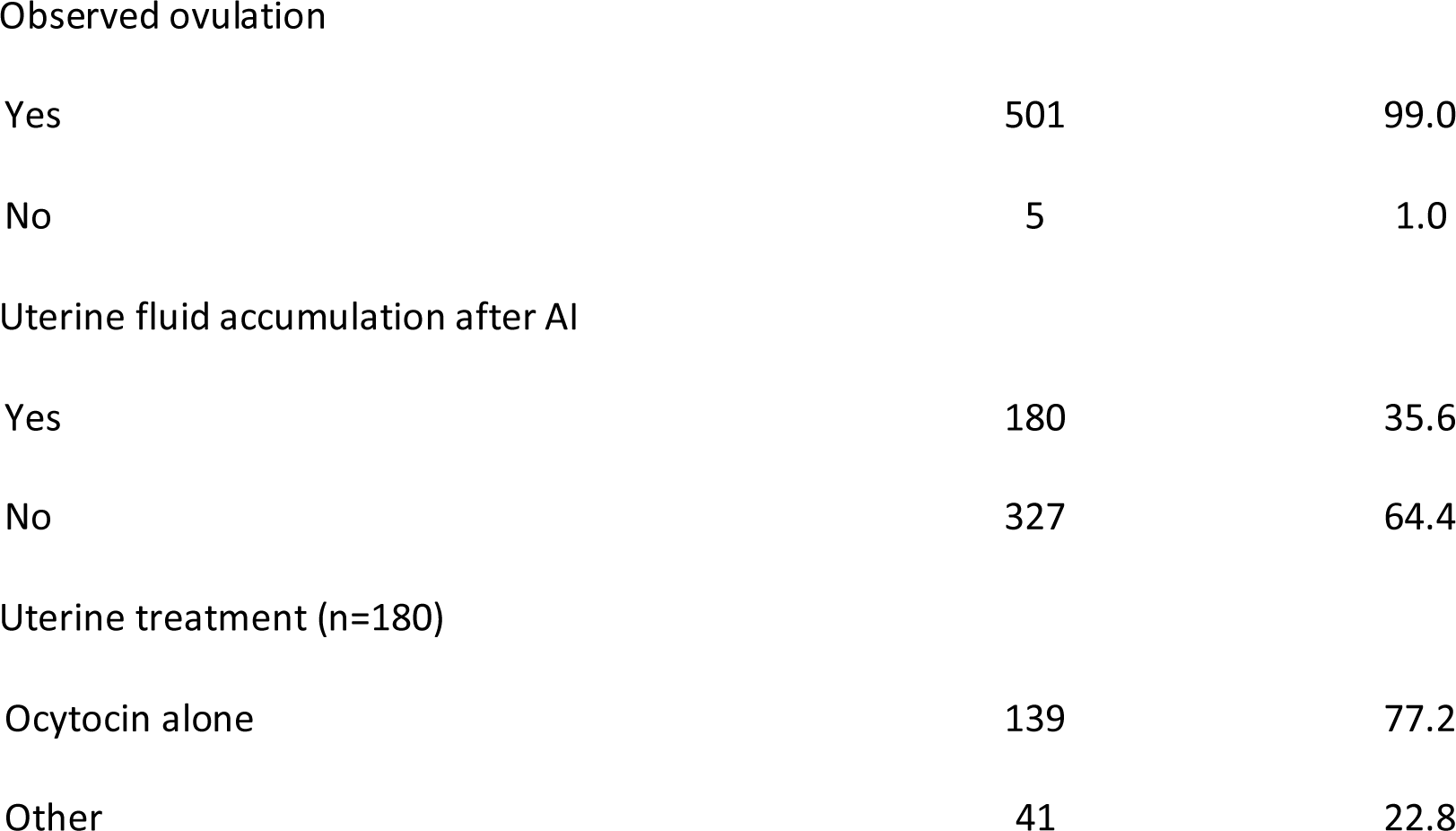
Characteristics of the 506 monitored estrus.

Among the 506 estrus recorded periods, 92 were induced by luteolysis of a previous corpus luteum using prostaglandins (18.2%). Data were recorded in average during 5.1 days per estrus period (min 1, max 16, σ = 2.1).

Of the 506 estrous periods, fresh semen was used for 90 cycles (FAI, 17.8%), cooled transported semen for 142 cycles (CTAI, 28.1%) and frozen semen for 274 (FZAI, 54.1%). Ovulation was induced in 360 estrous periods (71.2%). The induction of ovulation was performed in 40.0% of the FAI, 74.0% of the CTAI and 80.0% of the FZAI. Mostly hCG was used (n=322, 89.4%) with GnRH analog used in the other cases (n=38, 10.6%). Preovulatory dominant follicle mean diameter was 42.4 mm (min 28, max 60, σ = 4.8). Mares were inseminated in average 1.2 times per estrus (min 1, max 5, σ = 0.6). For FZAI, the mean number of straws used was 5.1 (min 1, max 12, σ = 2.4, recorded in 251 AI by 274).

Ovulation was observed by ultrasonography in 501 of the 506 estrous periods (99%). In most cases, only one ovulation was observed (n=425, 84.8%). Double or triple ovulations were observed in 75 (15.0%) and 1 estrus, respectively.

After insemination, uterine fluid accumulation was observed in 180 estrous periods (35.6%). Oxytocin was used as treatment for 173/180 cases, mostly alone (139/180), sometimes associated with uterine lavage (27 cases). Antibiotics were used for local treatment (10/180) associated to oxytocin (n=3), to uterine lavage (n=4) or alone (n=3).

After AI, 212 cycles among the 506 led to a pregnancy on D14 (pregnancy rate per cycle: 41.9%). Twins were detected in 27 cases (5% of breeding and 12.7% of pregnancies). Squeezing was performed for 25/27 pregnancies. After squeezing, pregnancy was checked on Day 30 and 21/24 mares were still pregnant (87.5%).

### 3.2. Univariate & multivariate analysis

#### 3.2.1. Factors associated with post-breeding inflammation

After univariate analysis, variables associated with post-breeding fluid accumulation were: mare’s age (using 10 years old as cutoff value), parity, lactation (suckling *vs* non suckling), induction of estrus by luteolysis, cycle number and month of AI (Figure 1). Post-breeding inflammation was not associated with semen conditioning (33.6, 40.2 and 37.9% respectively for FAI, CTAI and FZAI, p=0.45), number of AI per estrus (35.2% for 1 AI *vs* 40.3% for more than one AI per estrus, p=0.40), nor number of straws used (33.9 vs 33.8% for number of straws < 8 vs ≥ 8, p=0.98 and 37.3 vs 30.4% for number of straws ≤ 4 vs > 4, p=0.25).

**Figure 1:**
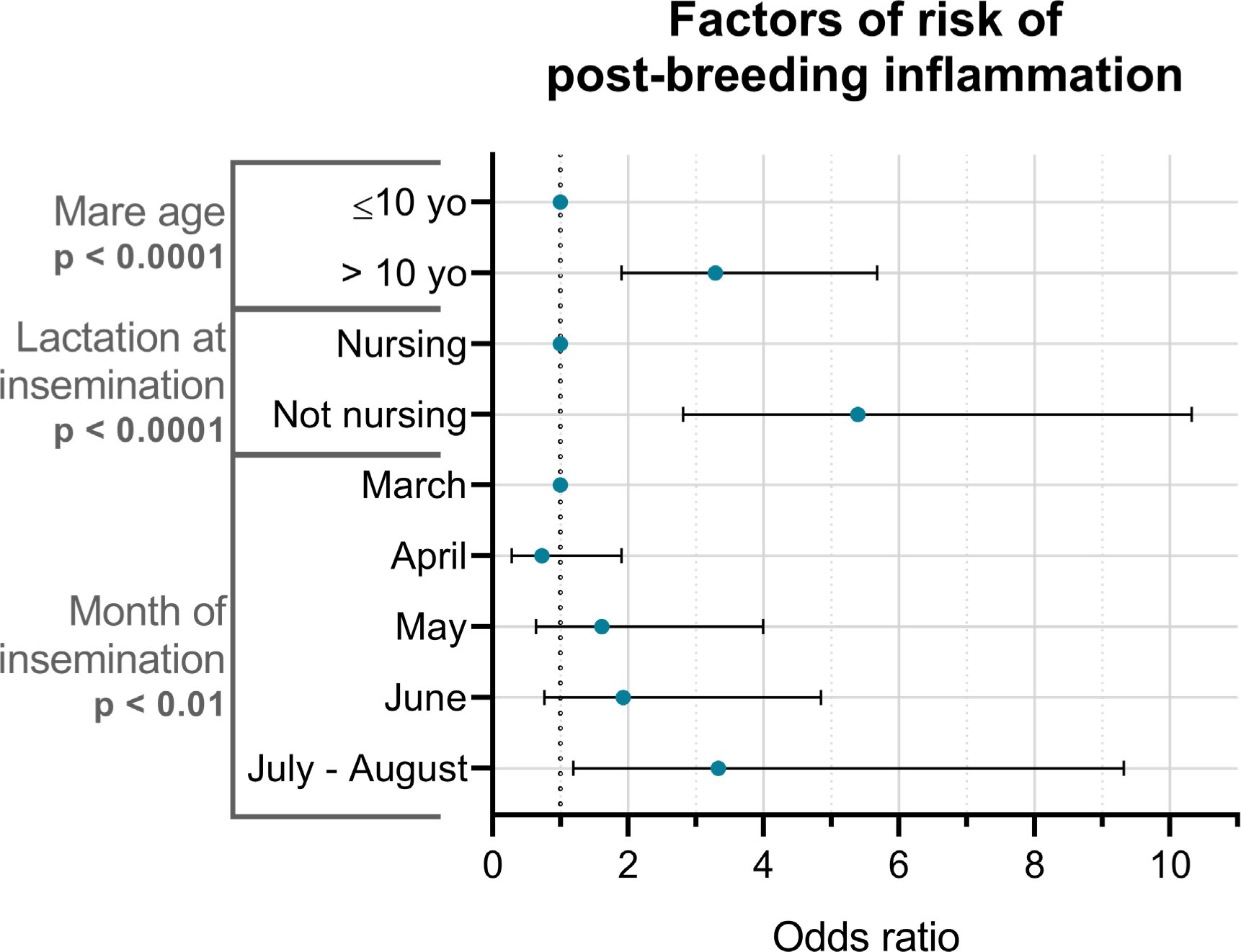
Odds ratio for factors influencing the likelihood of post-breeding endometritis in sport mares

After multivariate analysis, only 3 variables were significantly associated (p<0.05) with post-breeding inflammation: mare age, lactation and month of AI (Supplementary table 1). Inflammation was more frequent in mares older than 10 years than in younger mares (Figure 1). Non suckling mares at the time of insemination were also more affected in comparison with suckling ones (Figure 1). There was no interaction between age and nursing status (p = 0.38) but increased age amplified the number of post- breeding inflammations in both non-suckling and suckling mares. Respectively, 22.9% and 8.2% of the non-suckling and suckling young mares were affected by post-breeding inflammation while 50.7% and 18.3% of mares older than 10 years old were affected.

The risk of inflammation was also increased in July and August compared to previous months (Figure 1).

#### 3.2.2. Factors associated with pregnancy rate per cycle

After univariate analysis, the following factors were associated with pregnancy rate at the 20% threshold: age in 2 classes, parity, cycle number, AI modality, month of insemination, number of observed ovulations and post-breeding inflammation. All the associated variables were introduced in multivariate models. As a significant effect of number of ovulations was observed after univariate analysis, only estrous cycles with ovulations that were observed by ultrasonography were considered (*i.e.*, 501 estruses among the 506 recorded).

Stepwise regression and backward elimination led to a model containing only 3 significant variables (p<0.05): mare age, number of ovulations and semen conditioning (Supplementary Table 2). Data that significantly influenced pregnancy rates are summarized in Figure 2. Pregnancy rate was higher in mares younger *vs* older than 10 years old. Pregnancy rate was increased when multiple ovulations were observed. FAI and CTAI resulted in more pregnancies on Day 14 than FZAI. Trends were observed (p < 0.10) for the effects of month of AI and parity. Pregnancy rate tended to be higher in April, May, July and August vs March and June. Parous mares tended also to have better pregnancy rates than nulliparous mares (p=0.07, OR=1.434 in parous mares).

**Figure 2:**
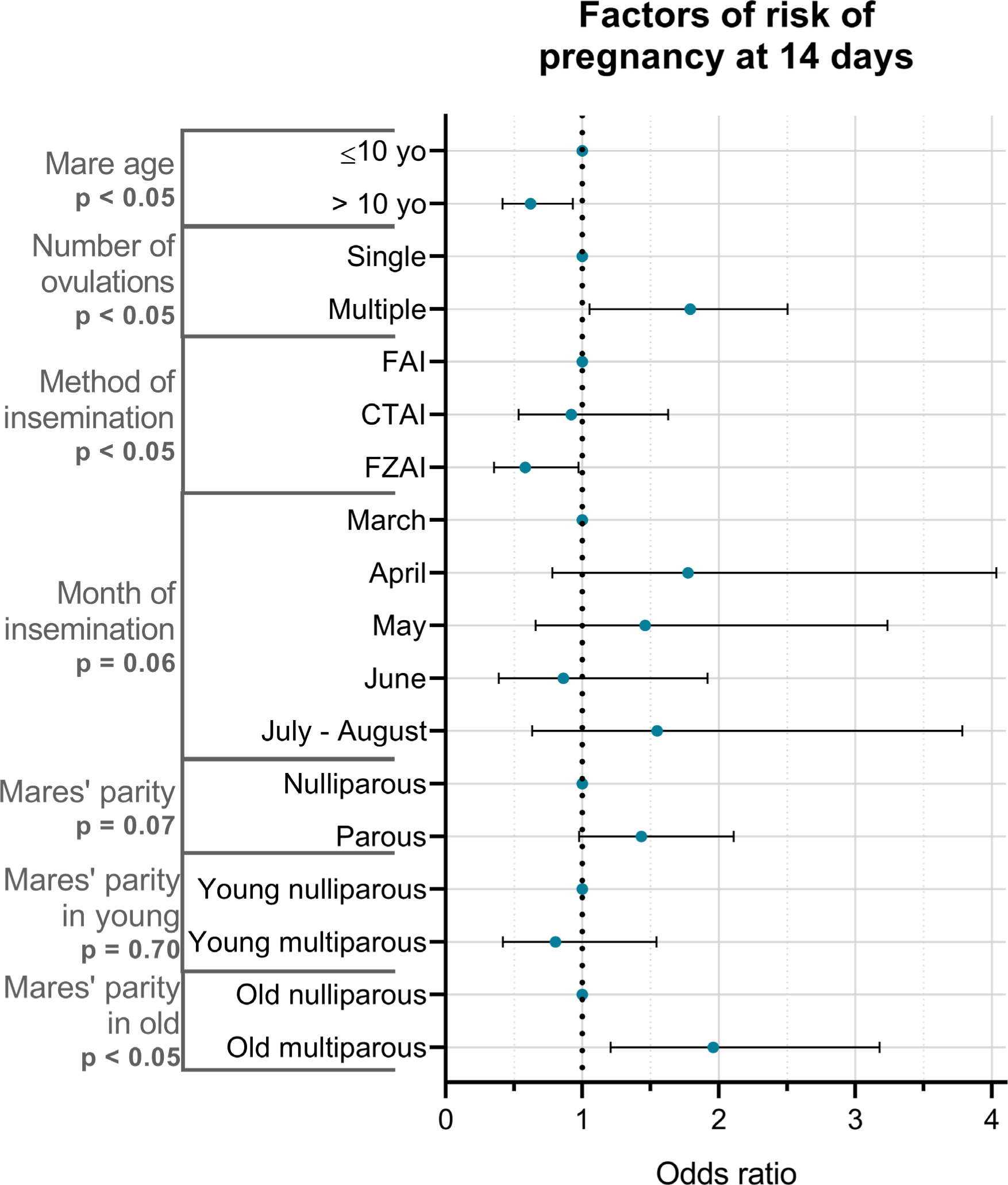
Odds ratio for factors influencing the likelihood of pregnancy in sport mares

The study of the interaction between maternal age and parity showed that in mares aged of 10 years or less, being nulliparous or parous did not alter pregnancy rates (49.25 and 46.15% of pregnancy rate per cycle for young nulliparous and parous respectively, n = 158, p = 0.70). In mares older than 10 years old, however, nulliparity accentuated the decrease in pregnancy rates. Indeed, the pregnancy rates per cycle was only 30.89% for old nulliparous mares *vs* 44.09% for old multiparous mares (OR = 1.96 in parous mares, p = 0.016).

After multivariate analysis, pregnancy rate was not significantly affected by post-breeding inflammation (37.8% of pregnancy rate after inflammation vs 44.2% in healthy mares, p = 0.16). In treated mares, treatment after post-breeding treatment did not affect pregnancy rate (37.4% of pregnancy for treatment with oxytocin alone, n = 139 *vs* 39.0% for other treatment, n = 41, p = 0.85).

#### 3.2.3. Factors associated with embryonic/fetal loss between Day 14 and foaling

After the univariate analysis, age in four classes, induction of ovulation and month of insemination were associated with embryonic/fetal loss at the threshold of 20%. However, after multivariate analysis, none of these factors affected embryonic/fetal loss.

### 3.3. Focus on maternal age, parity and suckling status

#### 3.3.1. Effect of maternal age

Mares 10 years old or younger were 7.2 ± 0.2 years old while mares older than 10 years were 15.6 ± 0.2 years old.

Neither parity, induction of the estrus cycle or of ovulation, month of AI, number of cycles required for a gestation at Day 14, number of straws in case of FZAI or the number of twin embryos at Day 14 were significantly related to maternal age (Supplementary Table 3).

Nevertheless, more young mares were pregnant at Day 14 and less had a post-breeding inflammation compared to old mares (Table 3). In addition, 31% of young mares were nursing while it represented only 20.7% of old mares (p < 0.05). Multiple ovulations occurred more frequently in old mares compared to young mares. Young mares tended to be more inseminated using fresh and frozen and less with refrigerated semen compared to older mares.

**Table 3:**
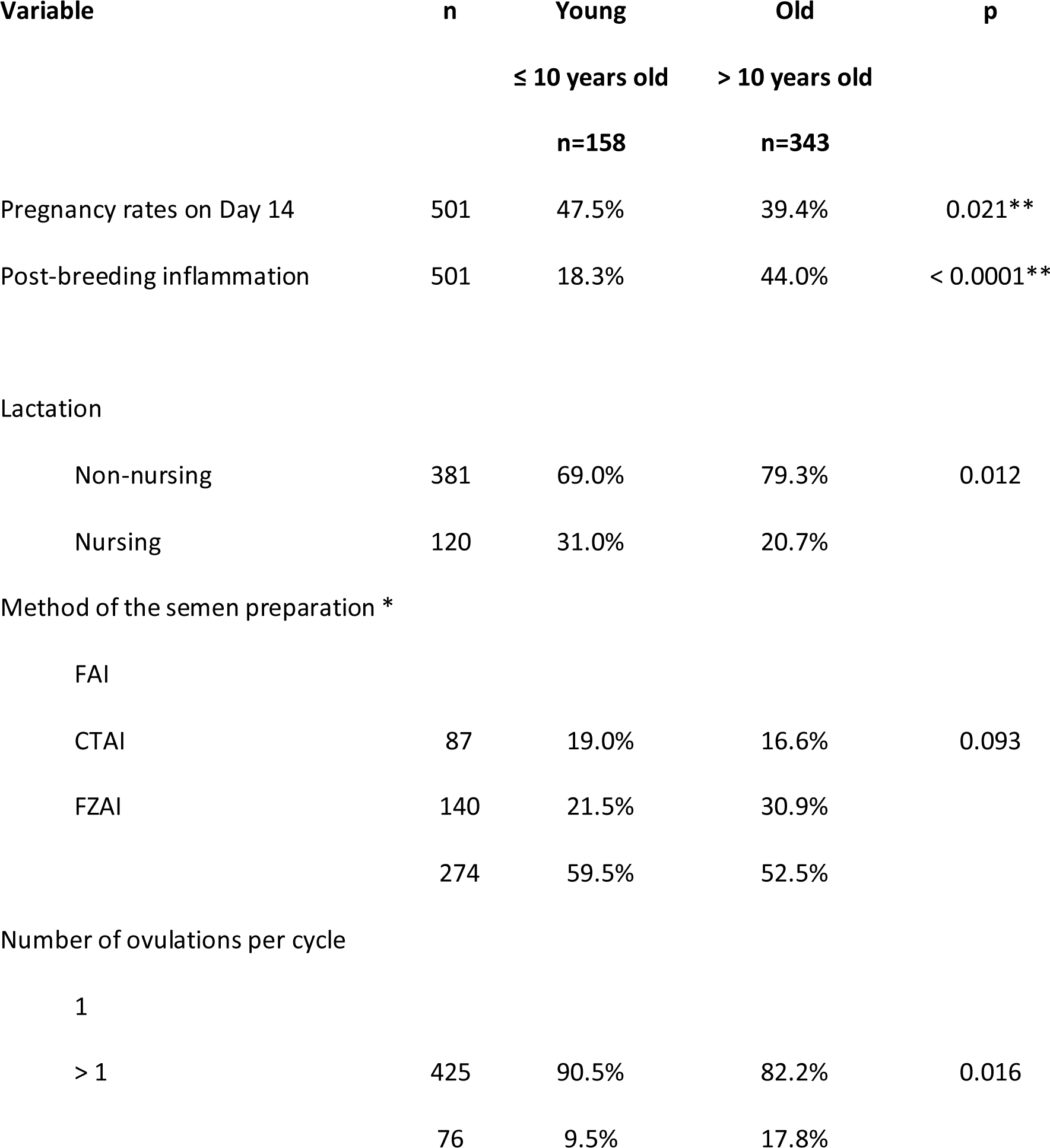
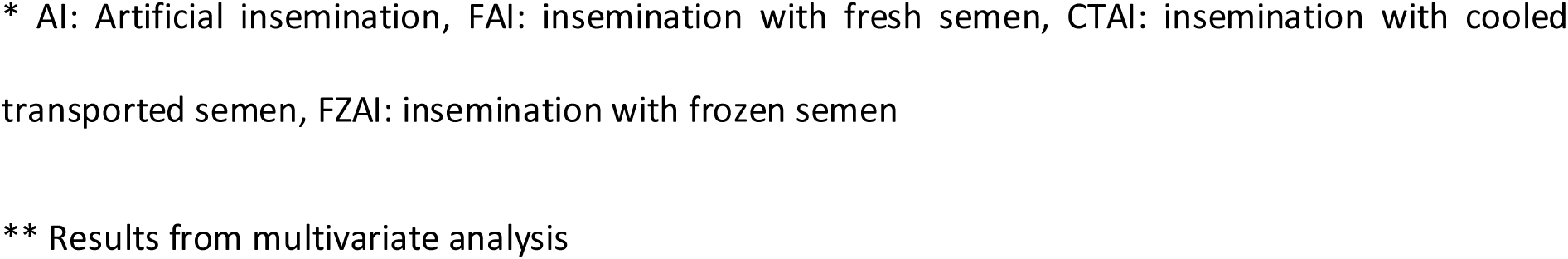
Characteristics in young (≤10 years old) vs old mares (>10 years old)

Estrus tended to be longer in old *vs* young mares (respectively, in average, 5.2 ± 0.1 *vs* 4.8 ± 0.2 days, p = 0.086). The preovulatory follicle diameter tended to be smaller in old compared to young mares (respectively, in average, 42.2 ± 0.2 *vs* 43.0 ± 0.4 mm, p = 0.086).

#### 3.3.2. Effect of maternal parity

Neither age, induction of the estrus cycle, semen preparation, month of AI, number of cycles or AI required for a gestation at Day 14, number of straws in case of FZAI nor the number of ovulation or twin embryos at Day 14 were significantly affected by maternal age (Supplementary Table 4).

Nulliparous mares tended to be less pregnant on Day 14 (Figure 2). Parity tended to be related to post-breeding inflammation. Maternal parity was different according to maternal age. More nulliparous mares were present in 5 or less and 10-15 years old group while less nulliparous were observed in 5- 10- and 16 or more-years old groups. Nulliparous mares were mostly older than 10 years (64.7%, Supplementary Table 4) but nulliparous were in average younger than parous mares (Table 4) with a mean 2-year difference between nulliparous and parous mares (in average, respectively, 11.6 ± 0.3 *vs* 13.7 ± 0.3 years old, p < 0.0001). Moreover, suckling status was obviously related to parity and 38.6% of parous mares were suckling at insemination. Ovulation was induced more frequently in parous than in nulliparous mares. Parity did not modify estrus duration (5.1 ± 0.2 days for nulliparous *vs* 5.1 ± 0.1 days for parous mares, p = 0.11) but the size of the preovulatory follicle at ovulation was reduced in nulliparous compared to parous mares (respectively, 41.9 ± 0.3 and 42.8 ± 0.2mm in diameter, p=0.03).

**Table 4:** Characteristics in nulliparous vs parous mares.

| Variable | n | Nulliparous | Parous | p |
| --- | --- | --- | --- | --- |
|  |  | n=190 | n=311 |  |
| Pregnancy rates on D14 | 501 | 37.4% | 44.7% | 0.066** |
| Age |  |  |  |  |
| ≤ 5 | 40 | 17.4% | 2.3% | < 0.0001 |
| 5 < Age ≤ 10 | 118 | 17.9% | 27.0% |  |
| 10 < Age ≤ 15 | 185 | 43.2% | 33.1% |  |
| Age > 15 | 158 | 21.6% | 37.6% |  |
| Lactation |  |  |  |  |
| Non nursing | 381 | 100.0% | 61.4% | < 0.0001 |
| Nursing | 120 |  | 38.6% |  |
| Induction of ovulation |  |  |  |  |
| No | 145 | 35.8% | 24.8% | 0.008 |
| Yes | 356 | 64.2% | 75.2% |  |

AI: Artificial insemination, FAI: insemination with fresh semen, CTAI: insemination with cooled transported semen, FZAI: insemination with frozen semen
\*\* Results from multivariate analysis

#### 3.3.3. Effect of nursing at insemination

Neither induction of the estrus cycle nor ovulation induction, semen preparation, number of AI per estrus required for a gestation at Day 14, number of straws in case of FZAI nor the number of ovulations or twin embryos at Day 14 were significantly affected by maternal suckling status (Supplementary Table 5).

Significantly less suckling mares were affected by post-breeding infection than non-suckling ones (Table 5) but pregnancy rates were not affecting by lactational status (45.8% and 40.7% of pregnancy rate per cycle for, respectively suckling and non-sucking mares, p = 0.32).

**Table 5:** Characteristics in non-suckling *vs* suckling mares.

| Variable | n | Non suckling<br>n=381 | Suckling<br>n=120 | p |
| --- | --- | --- | --- | --- |
| Post-breeding inflammation | 501 | 42.8% | 14.2% | < 0.0001 |
| Age |  |  |  |  |
| ≤ 10 years old | 158 | 28.6% | 40.8% | 0.012 |
| > 10 years old | 343 | 71.4% | 59.2% |  |
| ≤ 5 | 40 | 9.7% | 2.5% | < 0.0001 |
| 5 < Age ≤ 10 | 118 | 18.9% | 38.3% |  |
| 10 < Age ≤ 15 | 185 | 39.6% | 28.3% |  |
| Age > 15 | 158 | 31.8% | 30.8% |  |
| Month of AI |  |  |  |  |
| March | 35 | 8.1% | 3.3% | 0.005 |
| April | 103 | 22.8% | 13.3% |  |
| May | 149 | 29.7% | 30.0% |  |
| June | 153 | 26.7% | 42.5% |  |
| July - August | 61 | 12.6% | 10.8% |  |
| Number of cycles for pregnancy |  |  |  |  |
| 1 | 276 | 52.5% | 63.3% | 0.037 |
| > 1 | 225 | 47.5% | 36.7% |  |

\* AI: Artificial insemination, FAI: insemination with fresh semen, CTAI: insemination with cooled transported semen, FZAI: insemination with frozen semen

There were more suckling mares younger than 10 years of age and more non-suckling mares aged more than 10 years. Mares younger than 5 and mares aged between 10 to 15 years were less frequently nursing than mares aged 5-10. In the group of 5-10 years old, more suckling mares were present in comparison to non-suckling ones while the lactational status was similarly represented in 16 years old and older mares. In average, however, the age of suckling *vs* non-suckling mares was not different (respectively 13.0 ± 0.2 *vs* 12.6 ± 0.2 years old, p = 0.39). Less nursing mares were bred in March and April but more were bred in June than non-nursing ones (Table 5). Moreover, more nursing mares were pregnant at Day 14 within the first exploited cycle than non-nursing ones.

Estrus was shorter (respectively, 4.7 ± 0.16 *vs* 5.2 ± 0.1 days, p = 0.03) and preovulatory follicle diameter was larger (respectively 43.8 ± 0.4 *vs* 42.0 ± 0.2, p < 0.0001) in nursing vs non-nursing mares.

## 4. Discussion

Data presented here are summarized in Figure 3.

**Figure 3:**
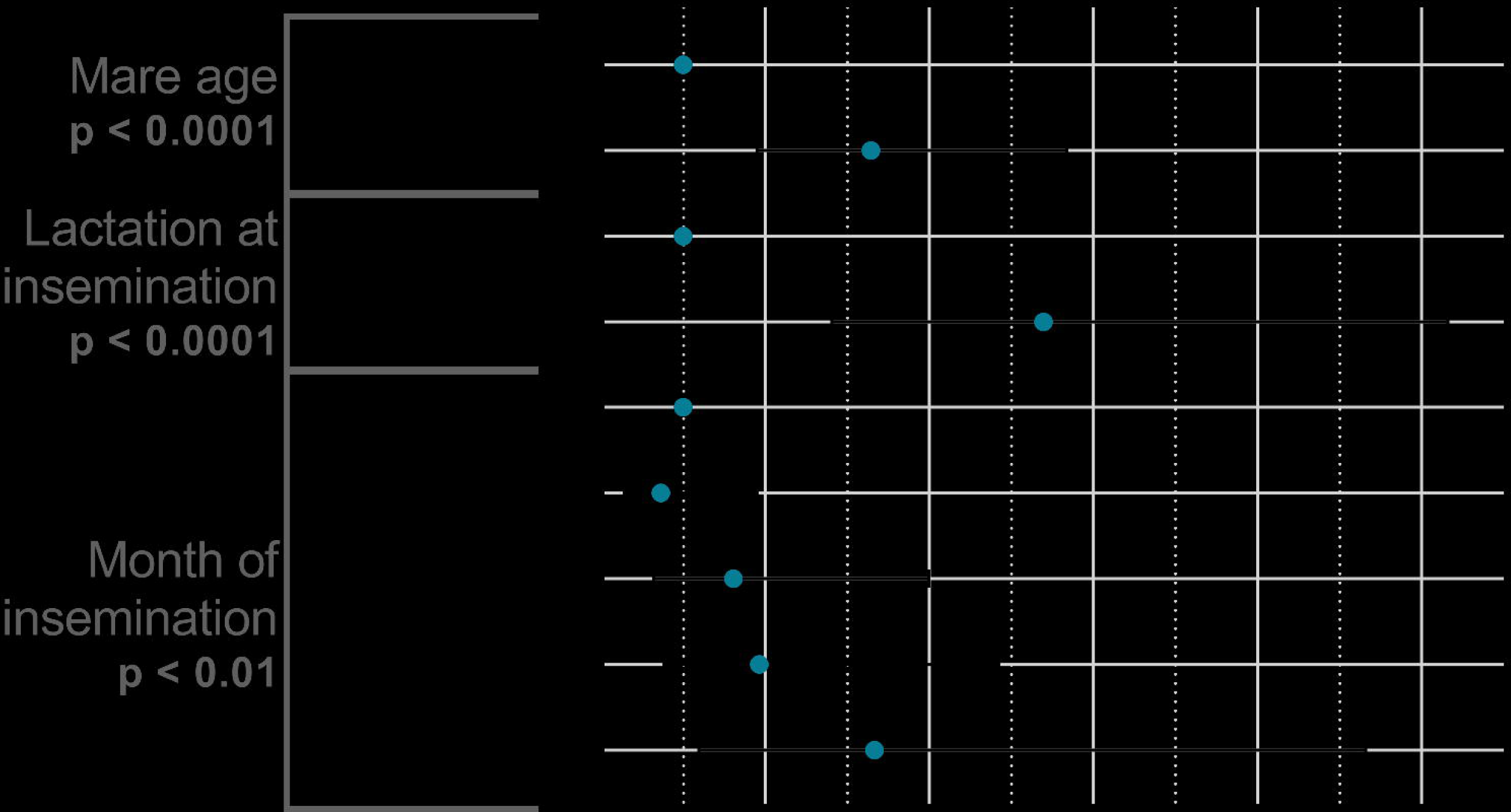
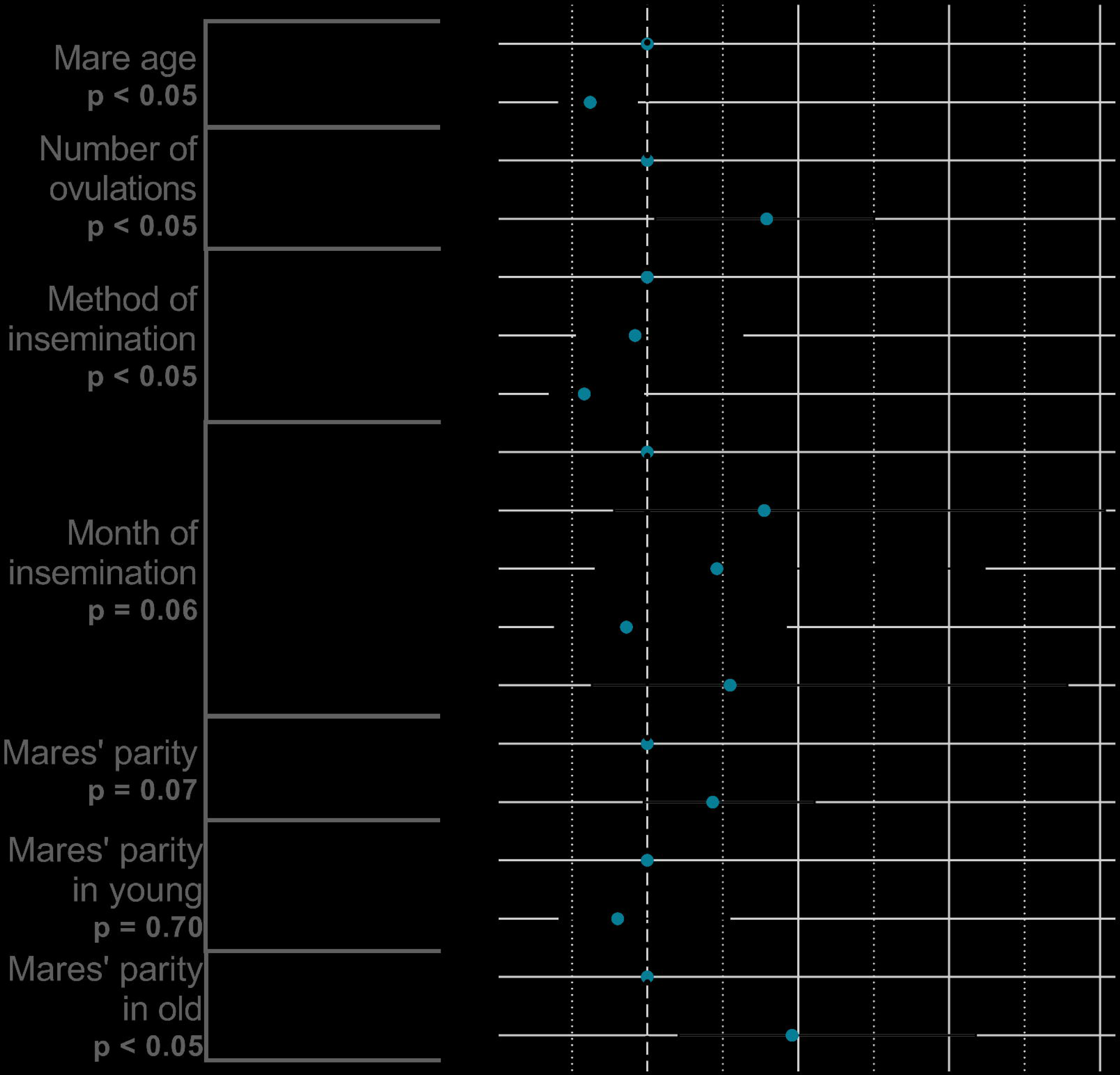
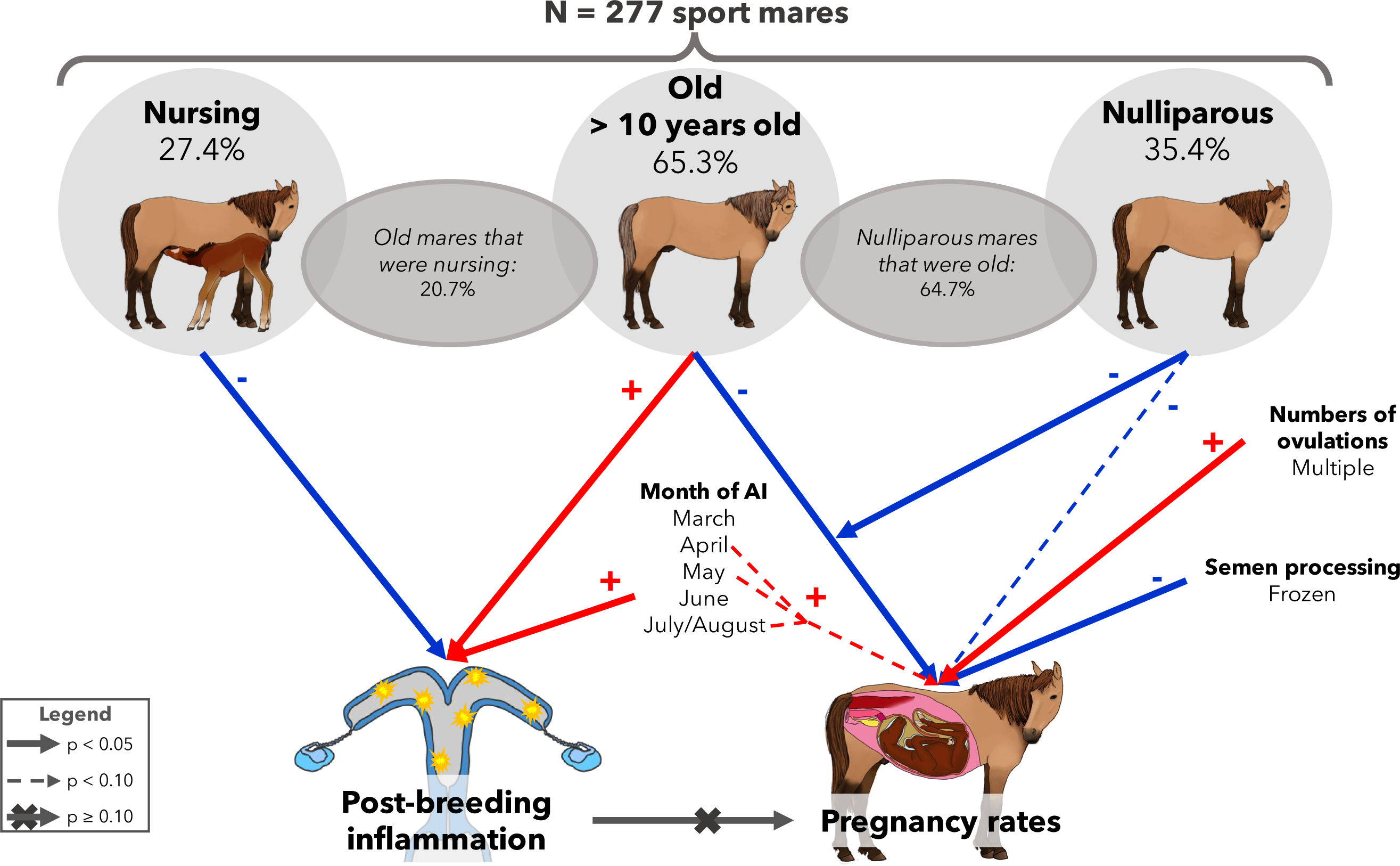
Factors affecting post-breeding endometritis and pregnancy rates in the studied population of sport mares Red arrows significates that the factor increased the likelihood while blue arrows indicated that the factor decreased the likelihood.

### 4.1. Post-breeding fluid accumulation

#### 4.1.1. Prevalence

In this study, more than a third of the monitored cycles were followed by post-breeding uterine fluid accumulation. In a normal population of Thoroughbred mares, around 15% of mares are susceptible to post-breeding inflammation that persists for several days [28]. In a more recent study in Thoroughbreds in UK, however, post-breading fluid accumulation occurred in 47.7% of analyzed pregnancies [17]. The observed rate in the present study is therefore in the range of the literature. In comparison to other studies, however, the post-breeding inflammation rate might have been overestimated. Indeed, as mares could be kept on the breeding stud for several cycles, the same mares could have had several inflammations over the entire breeding period, as this condition is generally persistent [29].

#### 4.1.2. Effect of maternal age, parity and lactation status

In the studied population, there was twice more post-breeding fluid accumulation in mares older than 10 years than in younger mares. These results agreed with literature as one recent study in Thoroughbred stud farms showed that the percentage of mares with post-breeding intrauterine fluid accumulation [17], increase with mare age. Frequency of endometritis has been demonstrated to increase with mare age as well as [26,30–34].

In addition, suckling at insemination appeared to have a protective effect against post-breeding fluid accumulation. To the authors’ knowledge, there is no study on the effect of nursing at insemination on post breeding fluid accumulation. Since lactating mares are producing endogenous oxytocin, the well-known and widespread treatment against post-breeding inflammation [35], it could explain the observed improvement in nursing mares. Indeed, nursing induces oxytocin release in the blood circulation, reaching a peak of around 10mIU/L in nursing pony and broodmares [36,37]. Plasma oxytocin concentrations of 10-7 mIU/L were reported to be present 20min after the beginning of the suckling period [36]. Since foals nurse until 72times a day in the first weeks of their life, the release of oxytocin happens in a regular manner throughout the day [38].

#### 4.1.3. Effect of the method of semen preservation and semen quantity

In this study, neither semen preservation method nor the volume determined by number of frozen straws, nor the number of inseminations were related to post-breeding fluid accumulation. One study showed a similar volume of uterine fluid accumulated in the uterus after insemination with cooled or frozen semen which is consistent with the results obtained here [39]. One recent study, however, showed that the uterine inflammatory response was positively corelated to the number of spermatozoids used for the insemination with frozen semen [40]. This high inflammatory response was also faster to resume than the response for low doses of semen [40].

#### 4.1.4. Effect of month of insemination

Data showed that more post-breeding fluid accumulation occurred when insemination was performed late in the season, i.e., during July and August. Several hypotheses could explain this result. The first is that mares with good quality uteri needed less cycles to become pregnant and therefore, more mares with reproductive troubles were inseminated late in the season as they often need to be inseminated for several cycles to start a pregnancy. Another hypothesis regarding the increase of post-breeding inflammation late in the season would be the increase of temperatures occurring from June/July . So far, there is no study on the effects of outdoor temperature on the incidence of endometritis in mares but in dairy cows, environmental heat, as evaluated by comparing the possibility of sheltering or not from the sun, increased rectal temperature by almost one degree [41] and reduced uterine blood flow [42]. The temperature increase, out of the thermoneutrality zone (5-25°C for horses), could directly affect uterine temperature and/or on uterine blood flow and therefore promote inflammation in the mares’ uterus.

#### 4.1.5. Treatment and pregnancy rates

In the present study, when a post-breeding fluid accumulation was detected, treatment was systematically applied. Most of the time, this treatment was limited to an injection of oxytocin. In a recent study about therapeutic practices in intensively managed Thoroughbred mares, almost half of the pregnant mares were treated with intrauterine antibiotics and the same proportion was treated with oxytocin. Oxytocin combined with intrauterine antibiotics are used prophylactically in Thoroughbreds in the UK to avoid uterine infections as breeders believe it to be a cause of conception failure and embryo loss [17]. In Thoroughbreds, however, less than 10% of early pregnancy failures were associated with uterine inflammation [17,31]. Another study on 99 Thoroughbred mares showed no association between post-breeding uterine fluid accumulation and pregnancy rates nor embryonic death [43], as observed here. In all studies, the systematic management to prevent/cure anormal inflammation could explain this lack of association. Therefore, nowadays, the post-breeding inflammation seems to be well handled by stud farms.

Contrarily, in different breeds, the combination of antibiotics with oxytocin to reduce post-breeding fluid accumulation was shown to be more efficient than antibiotics or oxytocin alone to prevent decreased pregnancy rates after mating [44]. In another study on warmblood mares artificially inseminated, however, oxytocin alone was sufficient to increase pregnancy rates in comparison to no treatment [35]. In this study, oxytocin alone was mostly applied and was sufficient to avoid altered pregnancy rate. Therefore, it suggests that this treatment alone is efficient to avoid decreased pregnancy rates in artificially inseminated mares.

### 4.2. Pregnancy rates

#### 4.2.1. Prevalence

In studies on different breeds conducted in several countries using artificial insemination, pregnancy rates were between 40-80%, according to the semen preservation method [15,45–47]. In a comparable recent study on 328 sport mares artificially inseminated in the Netherlands, 46.6% of gestations were obtained on Day 12 – 18 [48]. Here, pregnancy rates per cycle were similar with 41.9% of mares being pregnant by cycle regardless of the method used for semen preservation.

#### 4.2.2. Effect of maternal age, parity and lactation status

Pregnancy rates were reduced when mares were older than 10 years old at the time of insemination. Several studies already showed a reduction of pregnancy rates with increased maternal age [1,7,10,30,49–51]. Most studies, however, observed differences in mares older than 14 years old and not as early as 10 years old. Here, the clustering in 4 classes (≤ 5, 5-9, 10-14 and ≥ 15 years old) was not associated with alteration of pregnancy rates. Our inability to demonstrate such changes might be due to the only slight reduction of pregnancy rates induced by aging and to the limited number of mares studied.

Maternal parity tended to not reduce pregnancy rates in the overall population. The effects of maternal parity on pregnancy rates are contrasted as being closely related to mares’ age. Indeed, when nulliparous mares were mostly older than 7 years, a detrimental effect of nulliparity was observed [2,15] but when nulliparous mares were younger, no deleterious or a favorable effect of parity was observed [2,14]. Here, more than 60% of nulliparous were older than 10 years which could explain the observed tendency.

When considering only mares older than 10 years, D14 pregnancy rates were reduced in nulliparous mares while in the youngest group, the parity did not affect the incidence of pregnancy. Thus, the present data indicate a deleterious cumulative effect of nulliparity and aging on fertility in mares. This was previously suggested but never demonstrated in any broodmares population.

One other important finding is that lactation at the time of insemination did not influence pregnancy rates, 14 days post ovulation. Foaling mares, however, needed less cycles to become pregnant. In the literature, the effect of lactation on fertility is controversial as some studies observed that nursing mares are more fertile than non-nursing ones [2,8,30,50,51] while others do not observe any difference [4,7,10,14]. Insemination at foal heat was previously shown to reduce pregnancy rates [23,52] and to be associated with increased embryonic death [21,22,52]. Here, most foaling mares were bred on foal heat without adverse effect on pregnancy rates.

#### 4.2.3. Effect of the number of ovulations

The more ovulations were observed, the higher the likelihood that the mare was to be pregnant at 14 days post ovulation. The result obtained here seems obvious as double ovulations increase the number of oocytes that could be fertilized.

#### 4.2.4. Effect of the method of semen preservation and semen quantity

As previously observed [45,46,48,53], the modalities of insemination influenced the probability for a mare to be pregnant on Day 14. Indeed, pregnancy rate was higher after FAI or CTAI than after FZAI. It is well known that stallion semen quality decreases with cryopreservation, thus a higher critical number of mobile spermatozoa per dose is required to reach the same pregnancy rates than with fresh semen [53]. In this study, the number of straws was not related to pregnancy rate, as also shown by others [45]. In France, the concentration of spermatozoa/straw is standardized to 100 millions of spermatozoa per ml by the National French Institute. Thus, stallion coming from abroad had a different spermatozoa concentration, usually lower, that could also impact the results. Despite these considerations, the protocol involving ultrasonography every 6h and deep intra-uterine AI after observed ovulation seemed to be effective to reach acceptable pregnancy rates with 4 straws or less.

#### 4.2.5. Effect of month of insemination

The month of insemination tended to influence the pregnancy rate, with reduced incidence of pregnancies in March and June. In another French study, March was one of the more prolific month in terms of foal productivity per mare, all breed considered (Thoroughbred, Standardbred and sport bred) [2]. In racehorse breeding, it is common to use light to advance the breeding season [54] since foals born early in the year have an advantage when they are sold as yearling. In sport horse breeding under European latitudes, however, March is often the beginning of the breeding season. At this time, mares enter the spring transitional period and less mares are bred, which could explain the reduced pregnancy rates on this month. The other French study also reported a fertility decrease when mares were bred in June and after [2]. The June effect, here, is more complicated to explain as July and August did not affect pregnancy rates. During their stay, all mares were housed in individual stable with no access to fresh grass. They were fed with hay, thus a change in food quality in June could not explain the present results. Nevertheless, increasing ambient temperature could explain the differences in pregnancy rates as temperatures reached a maximum of 35.8°C in one stud farm in June 2019. In July and August, temperature were also high with more than 33°C as maximal temperature in the 2 studied stud farms. The lack of difference in July and August could be explained by the fact that only few mares were bred during this period and that more attention could have been given to these last mares.

### 4.3. Effects of age on other reproductive outcomes

A survey sent to breeders participating in competing events, showed that mare’s age was the least important consideration for the selection of mares for reproduction [55].

#### 4.3.1. Choice of semen preservation method

Maternal age did not influence the breeders’ choice concerning ovulation management but influenced the modality of AI. Indeed, less frozen and fresh semen were chosen for the insemination of older mares. On one side, frozen semen is less efficient for producing a foal, as confirmed by literature [53] and this study. Moreover, reduced fertility has been observed in old mares, both in literature [for review see 47] and in the present study. The combination of both factors could explain that older mares tended to be inseminated less with frozen semen as the financial risk might have been too costly for breeders. In addition, currently, equine breeders do not use more straws for old than for young mares, even if it has been shown that increasing straw number could improve fertility [53].

#### 4.3.2. Length of estrus and ovulation outcomes

This retrospective study shows that the length of estrus tended to increase with mare age (only 0.4 day more in old mares), with a smaller preovulatory follicle. This was already observed as a prolonged interovulatory interval was associated with a prolonged follicular growth [57–64] and to a reduction in follicular growth and preovulatory follicular size in older mares [65–68].

In the studied population, more multiple ovulations were also observed in older mares. Several studies reported that the incidence of multiple ovulations continuously increases until the mare reaches 20 years of age [63,68–70]. More multiple ovulations in older mares, however, is not associated with more twinning at 14 days post ovulation. Studies have shown that even if fertilization appears to be equal between young and old mares, early embryo mortality is increased in older mares [49,71] which could explain the absence in increase of twin embryos observed at 14 days in older mares compared to younger ones.

### 4.4. Effects of parity on other reproductive outcomes

Nulliparous mares had smaller preovulatory follicles. As explained above, the monitoring of mares was not performed more than twice a day: since follicles continuously grow until breakdown, this observed difference should be considered with caution.

### 4.5. Effects of suckling at insemination on other reproductive outcomes

In sport horses, mares are often bred and subsequently foal during the spring. Foaling mares are most of the time pregnant at the beginning of the spring. It is, therefore, not surprising to observe a peak of breeding in May and June in foaling mares while barren mare breedings were more spread throughout the reproductive season. Moreover, as it appears that less estrus cycles were required to obtain a pregnancy in lactating mares, it is not surprising either to not observed many lactating mares in August.

Finally, the heat period was shortened and the preovulatory follicle was larger when mares were in lactation without changing the number of ovulations. To the author knowledge, there is no study on the effect of lactation at insemination time on estrus and follicle parameters.

## 5. Conclusion

In conclusion, maternal age appeared to be the most important factor affecting both post-breeding inflammation and pregnancy rates. Both could be explained by the degenerative changes that are observed in older mares. Breeders, should, therefore, be encouraged to pay more attention to the age of their broodmares and either, breed earlier in their life or culling them earlier to avoid excessive costs. Older mares are also of less interest concerning genetics point of view [72] that should also be considered by breeders. Moreover, as frozen semen was associated with decreased pregnancy rates, the use of fresh or cooled semen for the insemination of old mares should be advised.

In this study, it has also been shown that nulliparity as the same time to aging affected pregnancy rates. For their first gestation, mares should not be more than 10 years old to increase chances of pregnancy. One advice to owners could be to breed the mares before starting their sportive career to produce their first foal as it will help for the later inseminations.

Suckling at insemination prevented post-breeding inflammation. The most probable hypothesis could be that sucking provokes oxytocin release that is acting as a natural treatment of uterine fluid accumulation.

Here, pregnancy rates were not affected by post-breeding inflammation, although uterine fluid accumulation were more frequent than in other studies, showing that the excessive fluid accumulation related to insemination is well handled and that oxytocin only as a treatment is efficient.

## Acknowledgements

The authors tanks V. Lehuraux, A Jugland, B. Brisset and P. Drevillon from the Stud of “Charmoy” (89) and Stud of “Les Bréviaires” (78) for the availability of the data and for their hospitality during the visits of their organizations. This research did not receive any specific grant from funding agencies in the public, commercial, or not-for-profit sectors. Many thanks to Juliette Auclair-Ronzaud for the reviewing of this article.

## Author contribution

CG generated data. BG formatted datasets and performed the analyses. ED, JAR and PCP wrote the draft. All authors read, revised, and approved the submitted manuscript.

